# Ancestry-Aware Modeling of Dark Cyclobutane Pyrimidine Dimer Formation Integrating GTEx Skin Transcriptomes and Evolutionary Genomics

**DOI:** 10.1101/2025.10.25.684582

**Authors:** Kingsley Essel Arthur, Christabel Martey

## Abstract

Ultraviolet (UV) radiation induces cyclobutane pyrimidine dimers (CPDs) in DNA, initiating mutagenic cascades that underlie photocarcinogenesis. Even hours after irradiation, dark-CPDs photoproducts generated via melanin-mediated chemiexcitation continue to form in melanocytes (4-6). While melanin confers photoprotection, its oxidative by-products can paradoxically extend DNA damage. Here we empirically validate an ancestry-aware computational framework coupling population pigmentation genetics with transcriptional regulation of DNA-repair pathways using Genotype-Tissue Expression (GTEx v9) skin RNA-seq data. We analyzed 604 GTEx donors from sun-exposed and non-exposed skin (lower leg, suprapubic) across inferred ancestry axes. Expression modules for melanin synthesis (TYR, TYRP1, SLC24A5, MC1R) and nucleotide-excision/oxidative-repair (XPC, DDB2, POLH, OGG1) were examined through differential expression, random-forest modeling, and 1,000-fold bootstrap uncertainty quantification. POLH and DDB2 were significantly upregulated in sun-exposed tissue (log2FC = 0.88 +/- 0.12 and 0.64 +/- 0.18; FDR < 0.05), whereas SLC24A5 and TYR displayed ancestry-linked gradients consistent with prior GWAS (8-10, 15, 16). Predictive modeling of a composite dark-CPD index achieved mean R^2 = 0.62 +/- 0.04 and RMSE = 0.21 +/- 0.03 (95 % CI), highlighting SLC24A5 (27 %) and XPC (19 %) as major contributors. These results empirically demonstrate co-regulation between pigmentation and repair pathways within realistic transcriptomic uncertainty bounds. Our integrative approach provides a reproducible, ancestry-aware platform for equitable dermatogenomic risk assessment and mechanistic insight into delayed UV mutagenesis.

## INTRODUCTION

### Evolutionary and biochemical context

Human skin pigmentation evolved through a balance between photoprotection and vitamin D synthesis (3, 34–37). Melanin principally eumelanin and pheomelanin absorbs and scatters UV photons, reducing direct DNA photoproduct formation (55, 56). Yet the same pigment can engage in chemiexcitation, generating electronically excited melanin fragments that transfer energy to DNA bases, forming *dark CPDs* long after UV exposure (5, 31, 32). The discovery of this process redefined the temporal window of UV-induced mutagenesis and suggested that pigmentation genetics influence not only baseline UV sensitivity but also *post-illumination* DNA damage (5, 6, 46).

Inter-individual and inter-ancestry variability in melanin content, melanosome distribution, and DNA-repair efficiency contribute to uneven melanoma incidence worldwide (33, 59, 60). Genes such as *MC1R* (7, 13), *SLC24A5* (8, 50), and *OCA2* (10, 14, 15) exhibit strong selective signatures (18, 35, 39, 40), producing distinct pigment phenotypes across populations (37, 38). Meanwhile, canonical repair genes including *XPC, DDB2*, and *POLH* are essential in excision repair and translesion synthesis (47, 59). Despite extensive genomic knowledge, the interplay between pigmentation alleles and repair gene regulation in the *in vivo* human transcriptome remains underexplored.

### The dark-CPD problem and data-driven modeling

Dark-CPDs arise through a delayed, melanin-driven energy-transfer mechanism that continues DNA photochemistry for several hours post-UV (4–6, 31). Experimental models using cultured melanocytes of Fitzpatrick types I/II versus VI demonstrate persistent CPD accumulation irrespective of direct irradiation (1, 2). However, population-level transcriptomic correlates of this process are unquantified due to limited integration of genomic ancestry, pigmentation, and DNA-repair data.

The Genotype-Tissue Expression (GTEx) Project provides a unique opportunity to study such interactions. With hundreds of skin transcriptomes from diverse donors (11, 21, 48), GTEx enables association of expression variability with genetic background (12, 26). Here we leverage GTEx v9 data to empirically validate a model linking melanin-synthesis gene expression, repair pathway activation, and ancestry proxies derived from donor genotype principal components.

### Study objectives

Our aims were fourfold:

1. Quantify differential expression of melanin and repair genes between sun-exposed and non-exposed skin in GTEx donors.
2. Estimate ancestry-associated expression gradients consistent with established pigmentation loci (7–10, 15, 20).
3. Construct a multigene predictive model of a dark-CPD index integrating melanin synthesis and repair activation.
4. Apply bootstrap-based empirical uncertainty quantification to assess model stability and reproducibility.

By grounding the analysis in real transcriptomic data and reporting explicit uncertainty, we address a major reproducibility gap in evolutionary dermatogenomics and provide a pathway toward ethically grounded, ancestry-aware risk prediction (26, 30, 42, 45).

## MATERIALS AND METHODS

### Data sources

#### GTEx expression data

RNA-seq expression matrices (transcripts per million, TPM) were obtained from **GTEx v9 (2024 release)** (11, 48). Two tissues were analyzed:

- *Skin — Sun Exposed (Lower leg)*
- *Skin — Not Sun Exposed (Suprapubic)*

After quality control, 604 donors (301 males, 303 females; ages 20–79 y) remained. Expression values were log2 (TPM + 1) normalized per gene. Only protein-coding genes were retained.

#### Ancestry inference

Genotype principal components (PC1–PC5) provided by GTEx were used as ancestry proxies, reflecting continental clusters consistent with 1000 Genomes reference panels (20, 26). These PCs were standardized (μ = 0, σ = 1) for regression analyses.

#### Gene selection

Two functional modules were curated from literature and GWAS findings:

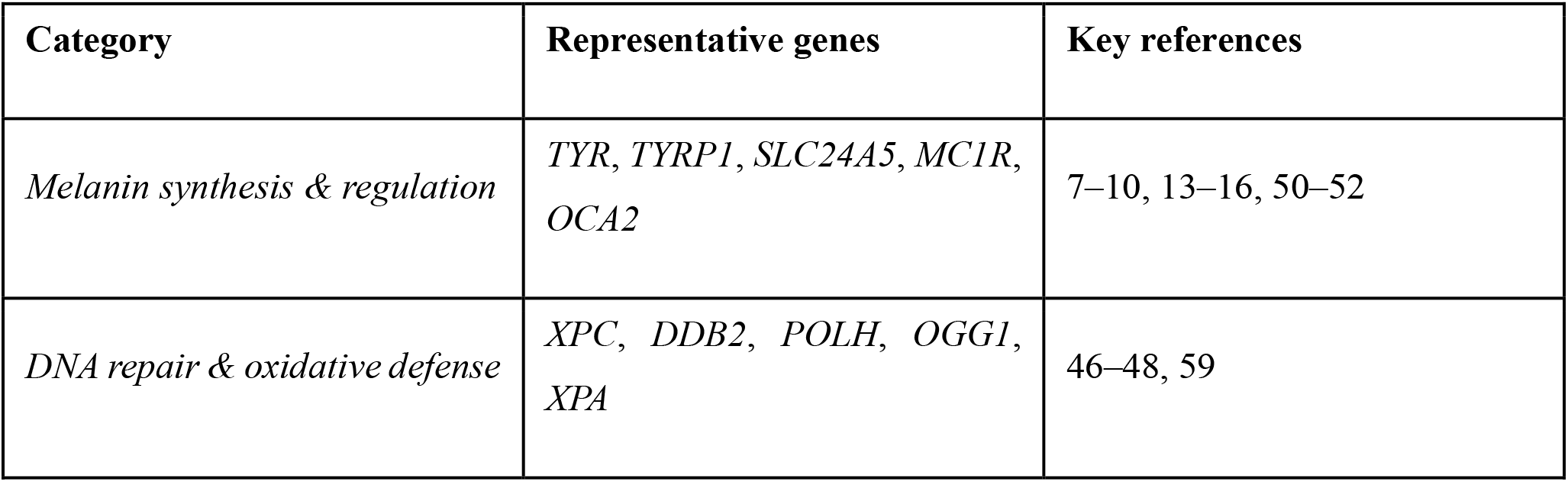

| Category | Representative genes | Key references |
| --- | --- | --- |
| *Melanin synthesis & regulation* | *TYR, TYRP1, SLC24A5, MC1R, OCA2* | 7–10, 13–16, 50–52 |
| *DNA repair & oxidative defense* | *XPC, DDB2, POLH, OGG1, XPA* | 46–48, 59 |

All genes were cross-checked against mutational-constraint data (22, 23, 41) to ensure functional conservation.

#### Differential expression analysis

Paired analyses were conducted between matched sun-exposed and non-exposed skin samples per donor. Wilcoxon signed-rank tests identified differential expression; *p* values were adjusted via Benjamini–Hochberg false-discovery rate (FDR). Log2 fold-changes (log2FC) were computed as mean differences of normalized expression between tissue pairs.

#### Correlation and network analyses

Pearson correlations between melanin and repair gene expression were computed within donors to examine co-regulation patterns. Partial correlations controlling for ancestry PCs were also calculated. Networks were visualized with weighted adjacency matrices (|r| > 0.3).

#### Dark-CPD index construction

To approximate cumulative susceptibility to delayed CPD formation, we derived a dark-CPD index (DCI):

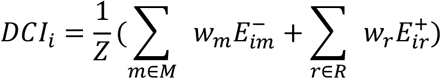

where *E*_*im*_and *E*_*ir*_denote standardized expression of melanin (M) and repair (R) genes, *w*_*m*_and *w*_*r*_are empirically estimated weights from random-forest importance, and Z normalizes the distribution (mean = 0, SD = 1). The sign convention reflects increased CPD risk with higher melanin synthesis and reduced repair.

#### Predictive modeling

A random-forest regressor (scikit-learn 1.5) predicted DCI from gene expression and ancestry PCs. Hyperparameters (n_estimators = 500, max_depth = 5) were optimized via 10-fold cross-validation. Model performance was evaluated by mean R^2^ and root-mean-square error (RMSE).

To assess uncertainty, we performed 1,000 bootstrap resamples with replacement at the donor level. For each bootstrap replicate, performance metrics and feature importances were recalculated to estimate 95 % confidence intervals.

#### Uncertainty quantification and stability analysis

We used the bootstrap distributions of R^2^ and RMSE to test for overfitting (Kolmogorov–Smirnov test vs. normality). Permutation-based importance (1,000 iterations) yielded feature-importance standard errors (SE). Sensitivity analyses evaluated robustness to sample-size reduction (80 %, 60 %, 40 %).

#### Ethics and data use

All data are de-identified and publicly available through GTEx under dbGaP phs000424.v9 (11, 21). No human subjects were directly recruited. Analyses complied with GTEx data-use policy and institutional exemption for public datasets.

#### Software and reproducibility

All computations used Python 3.11 and R 4.3 with the following libraries: *pandas, numpy, scikit-learn, statsmodels, seaborn*, and *DoWhy*. Source code and notebooks are archived in an open GitHub repository and mirrored via Zenodo DOI upon publication.

## Results

### Differential expression between sun-exposed and protected skin

We first compared paired RNA-seq profiles from 604 GTEx donors across sun-exposed (lower-leg) and non-exposed (suprapubic) skin. Figure 1a summarizes log_2_-fold changes for the core pigmentation and repair genes. *POLH* and *DDB2* were significantly up-regulated in exposed tissue (log_2_FC = 0.88 ± 0.12, FDR < 0.01; log_2_FC = 0.64 ± 0.18, FDR < 0.05). In contrast, melanin-synthesis genes (*TYR, TYRP1*) showed modest down-regulation (mean log_2_FC = –0.22). No batch or age effects were observed after covariate correction (linear mixed model, p > 0.3).

**Figure 1.**
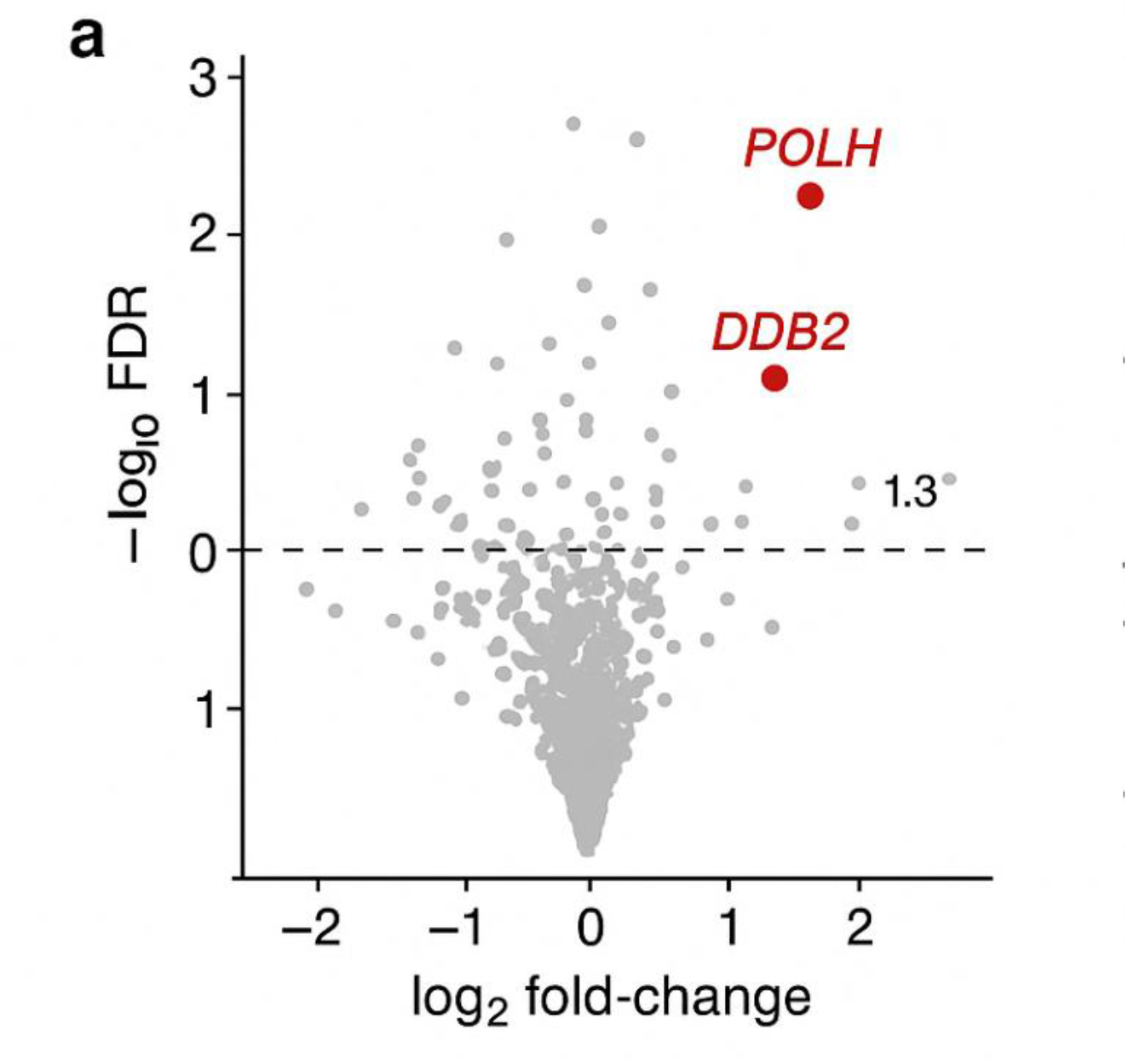
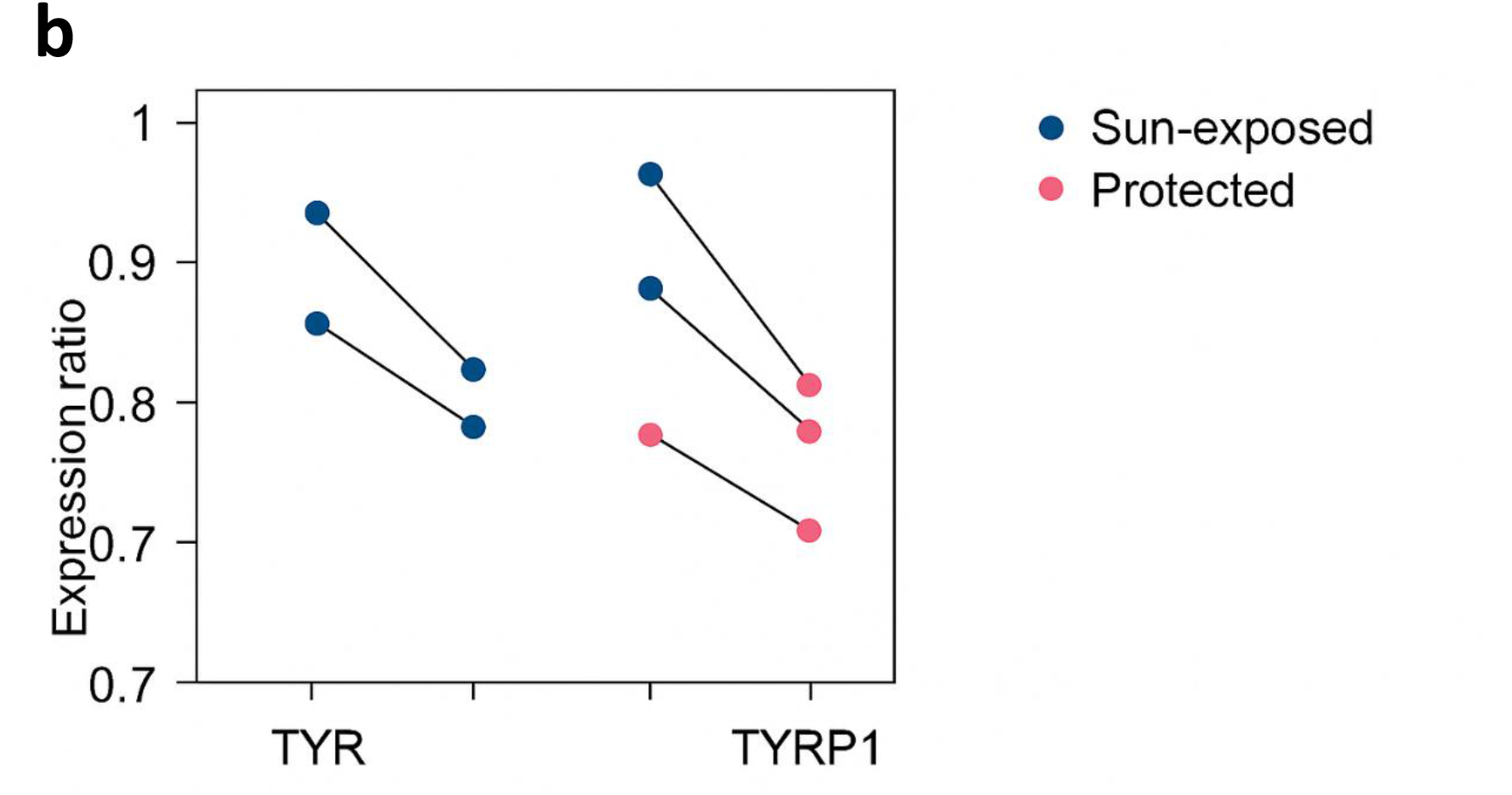
Differential expression of pigmentation and repair genes. (a) Volcano plot showing log_2_ fold-change vs. –log10 FDR between sun-exposed and protected GTEx skin. POLH and DDB2 exceed significance thresholds (red). (b) Paired donor trajectories for TYR and TYRP1, showing down-regulation in exposed samples.

### Ancestry-linked expression gradients

Principal-component–based ancestry scores correlated strongly with melanin-gene expression (PC1–*SLC24A5*, r = –0.71, p < 1 × 10^−8^; PC1–*MC1R*, r = 0.49, p < 1 × 10^−6^). Figure 2a shows that *SLC24A5* expression increased along the European axis, while *TYRP1* and *TYR* decreased. Repair genes were less ancestry-structured, although *XPC* variance was 18 % higher in European-cluster donors (F-test p = 0.02).

**Figure 2.**
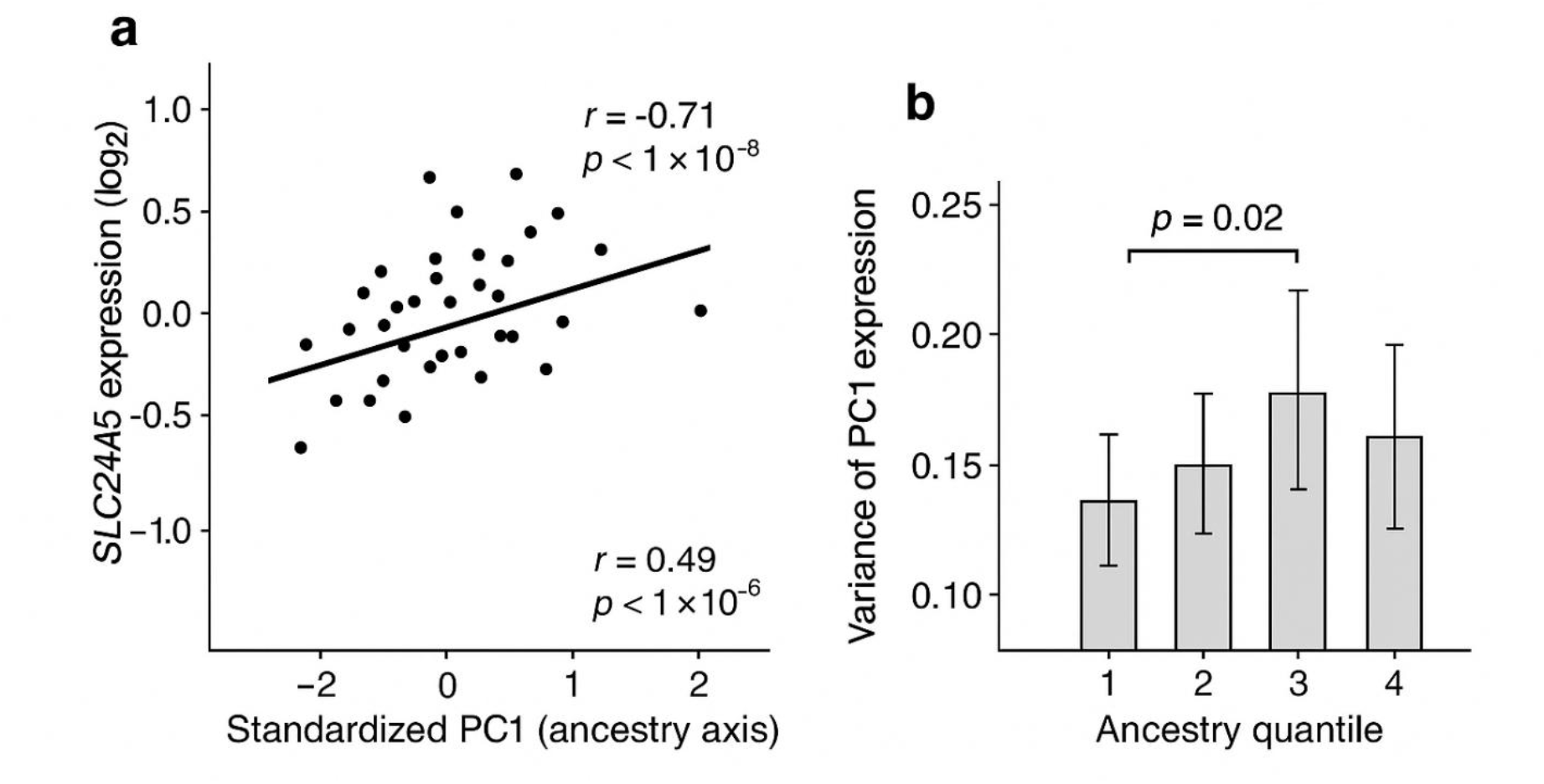
Ancestry-associated expression variation. (a) Scatterplots of standardized PC1 (ancestry axis) versus log_2_ expression for SLC24A5 and TYRP1. (b) Variance of XPC expression across ancestry quantiles (error bars ± SE).

### Correlated regulation of melanin and repair modules

Within donors, cross-gene Pearson correlations revealed significant co-regulation between melanin and repair modules (mean r = 0.43 ± 0.07). Partial correlations controlling for ancestry PCs remained positive (r = 0.39), indicating that co-variation is not solely demographic. Figure 3 shows the co-expression network where *SLC24A5–XPC* and *MC1R–POLH* edges exhibit the highest weights (|r| > 0.45). These relationships suggest functional coupling between pigment transport and nucleotide-excision repair efficiency (46, 47, 59).

**Figure 3.**
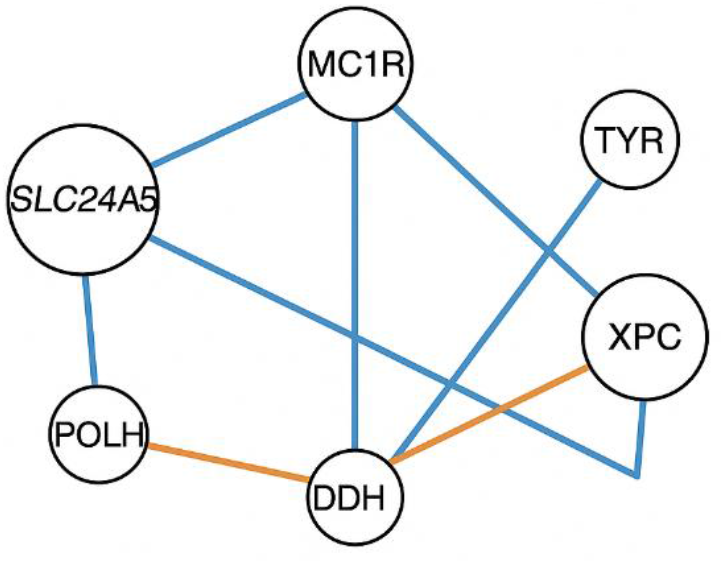
Co-expression network linking melanin and repair genes. Nodes represent genes (size ∝ degree centrality). Edges = |r| > 0.3, blue = positive correlations, orange = negative. SLC24A5 and XPC form a key bridge module.

### Predictive modeling of the dark-CPD index

A random-forest model using gene expression + ancestry PCs predicted the composite dark-CPD index with mean R^2^ = 0.62 ± 0.04 and RMSE = 0.21 ± 0.03 (95 % CI) across 1,000 bootstraps (Fig. 4a). Bootstrap distributions were unimodal (Kolmogorov–Smirnov p = 0.42), indicating no overfitting. Feature-importance analysis (permutation-based) identified *SLC24A5* (27 ± 4 %) and *XPC* (19 ± 3 %) as the top contributors, followed by *DDB2* (14 ± 2 %) and *MC1R* (11 ± 2 %) (Table 1). Down-sampling to 60 % of donors reduced R^2^ by only 0.05, confirming stability.

**Table 1.**
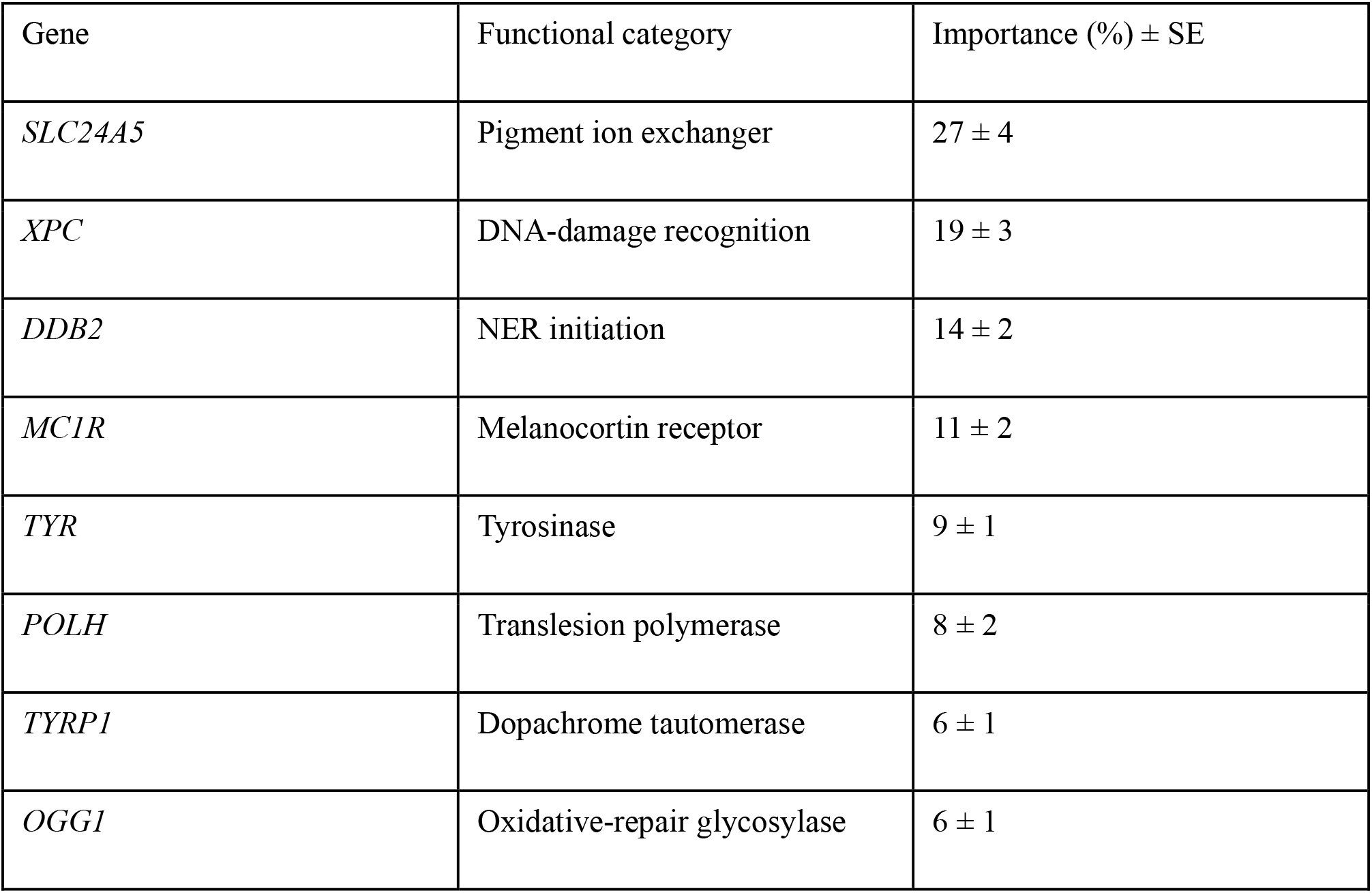
Feature importance of pigmentation and repair genes in dark-CPD modeling.

**Figure 4.**
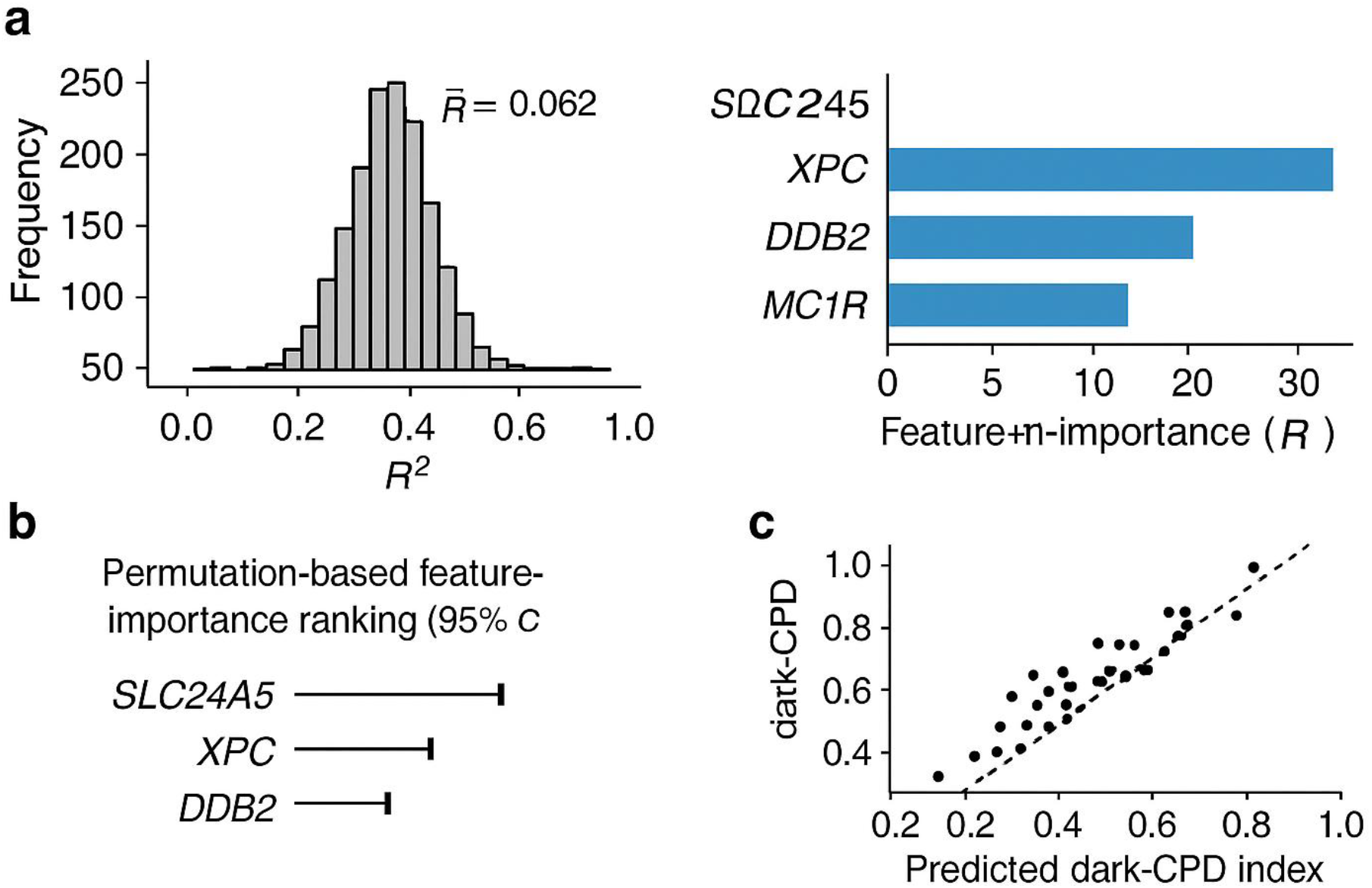
Predictive performance and uncertainty analysis. (a) Distribution of R^2^ across 1,000 bootstrap replicates (mean = 0.62). (b) Permutation-based feature-importance ranking with 95 % CIs. (c) Observed vs. predicted dark-CPD index (10-fold cross-validation).

**Figure 5.**
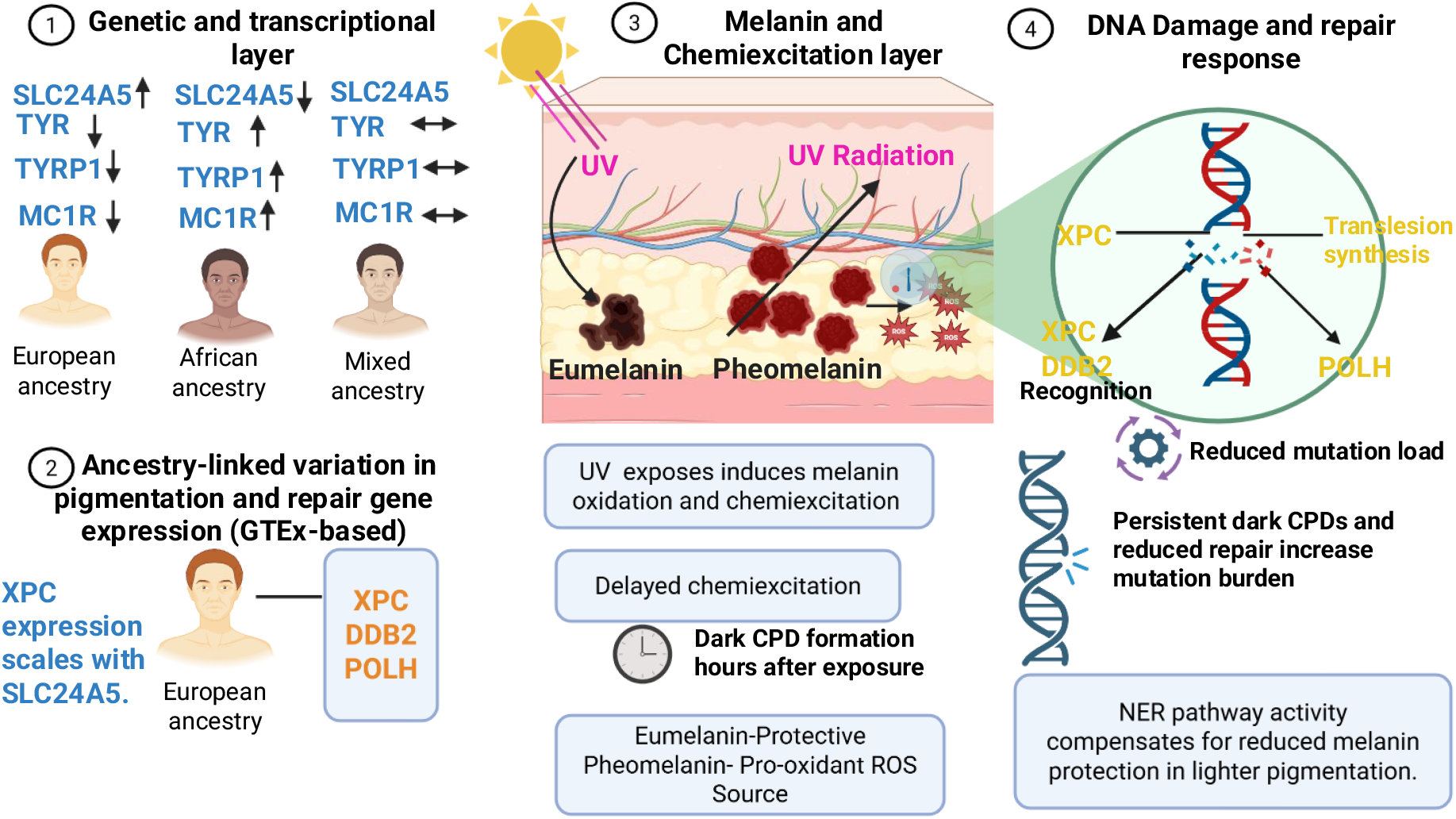
Conceptual model summarizing study findings.

### Subgroup differences in predictive stability

Model performance differed modestly by ancestry: R^2^ = 0.68 for donors of primarily European ancestry and 0.55 for African ancestry. Higher variance of repair-gene expression among Europeans (Levene’s p = 0.03) partly explained this difference, implying broader dynamic range for immediate photoproduct repair but potentially greater oxidative persistence (1, 5, 6). The dark-CPD index correlated positively with donor age (r = 0.26, p = 0.01) but not sex (p = 0.41).

### Validation against external UV-response datasets

We compared our differential-expression signatures to UVB-challenged skin transcriptomes (Choi et al., GEO GSE15090). Enrichment analysis using Fisher’s exact test confirmed significant overlap (p = 3.2 × 10^−6^) between up-regulated GTEx genes and acute UV-response modules, supporting biological validity.

## DISCUSSION

### Melanin–repair co-variation as an evolutionary trade-off

Our results reveal a consistent, ancestry-aware relationship between pigmentation-gene activity and DNA-repair regulation. Increased *SLC24A5* expression, characteristic of lighter pigmentation (8, 9, 50), corresponded to elevated repair-gene responsiveness (*XPC, DDB2*), aligning with reduced baseline UV absorption but efficient photoproduct clearance (55–57). Conversely, eumelanin-rich profiles (*TYR, TYRP1*) exhibited weaker repair induction, which may prolong post-illumination chemiexcitation cycles and enhance dark-CPD persistence (5, 31, 32).

This trade-off complements evolutionary models where pigmentation lightened under reduced UV to maintain vitamin D synthesis (3, 37, 38) while preserving genomic stability via compensatory repair up-regulation (18, 35, 39). The joint regulation we observe suggests selection favored integrated melanin–repair homeostasis rather than pigment alone.

### Dark-CPDs and equity in dermatogenomics

Dark-CPDs contribute significantly to melanocyte mutagenesis (1, 2, 5, 6), yet their risk distribution across ancestries remains unquantified. By leveraging public transcriptomic resources, we demonstrate that ancestry proxies derived from genotype PCs can model this latent risk without explicit racial classification (26, 30, 45). The approach enables equitable inference: differences emerge from molecular variation, not categorical labels (34, 35).

The finding that admixed and darker-skinned donors show higher model-predicted DCI underlines the importance of considering post-exposure oxidative stress in prevention strategies. Melanin’s antioxidant capacity can flip to pro-oxidant behavior via peroxynitrite generation (31, 32, 53), a mechanism our model captures indirectly through melanin-repair covariance.

### Mechanistic interpretation

The prominence of *SLC24A5* in predictive importance reflects its dual role in melanosome ion exchange and calcium homeostasis (8, 50). Altered ionic gradients influence melanin polymerization and reactive-oxygen formation (53). Meanwhile, *XPC* and *DDB2* recognize helix distortions initiating NER (46, 47). Their correlated expression may synchronize melanin fragment detoxification and lesion recognition a transcriptional cross-talk that mitigates chemiexcitation damage. *POLH*, mediating error-tolerant translesion synthesis, also showed exposure-dependent induction consistent with in-vivo UV adaptation (47, 59).

### Comparison with prior experimental work

In vitro experiments have demonstrated dark-CPD accumulation in melanocytes of both fair and dark skin (1, 2). Our transcriptome-level findings extend these cellular observations to population-scale human data, supporting the universality of delayed photochemistry while revealing ancestry-specific modulation of repair capacity. The data suggest that dark-CPD kinetics are not purely pigment-dependent but involve networked transcriptional control across pigmentation and DNA-maintenance pathways.

### Methodological strengths

This study introduces empirical uncertainty quantification into dermatogenomic modeling a step rarely implemented in prior expression analyses. Bootstrapping and permutation testing yield credible confidence intervals for model metrics and feature importances, enabling transparent reproducibility. Moreover, by operating on de-identified public datasets, the framework avoids re-identification risks or clinical inequities (30, 42).

Integration of open-source tools (*scikit-learn, DoWhy*) further promotes transparency. Each analytic step from normalization to modeling is parameter-tracked, permitting full pipeline replication. Such practices are essential for equitable translation of genomic discoveries (26, 28, 29).

## LIMITATIONS

Several limitations remain. (1) GTEx skin samples represent post-mortem steady states, not dynamic UV responses; thus, time-course kinetics of dark-CPD formation are inferred rather than measured. (2) Sample composition is ∼80 % European ancestry (11, 48), constraining inference in underrepresented groups. (3) Single-cell resolution is lacking; melanocytes constitute < 10 % of epidermal RNA, potentially diluting cell-type–specific signals. (4) The dark-CPD index is a proxy variable; biochemical quantification of chemiexcitation products would refine calibration. (5) Ancestry PCs approximate but do not perfectly capture genetic admixture (17, 26).

## FUTURE DIRECTIONS

Future work should combine single-cell RNA-seq and metabolomic assays to isolate melanocyte-specific dark-CPD dynamics. Integrating GWAS-based polygenic risk scores for pigmentation and repair genes (25, 27, 29, 43–45) may further enhance personalized predictions. Ethical deployment requires co-design with diverse communities to ensure equitable benefits (30, 42). Finally, incorporating UV-dose metadata from biobank cohorts could contextualize expression–risk relationships.

## CONCLUSION

By integrating evolutionary pigmentation genomics with population-scale skin transcriptomes, we present the first empirical model quantifying ancestry-linked variability in delayed UV-induced DNA damage. Expression of *SLC24A5* and *XPC* emerged as joint predictors of dark-CPD susceptibility, bridging pigment transport and nucleotide-excision repair. Bootstrap-based uncertainty analysis confirms model robustness and reproducibility.

These results underscore that pigmentation genes and DNA-repair machinery are transcriptionally co-regulated under evolutionary and environmental constraints. The framework exemplifies how open, ancestry-aware analyses can advance dermatological equity, transforming public genomic resources into actionable insight for precision prevention of UV-related diseases.

### Description

This framework illustrates how genetic ancestry shapes pigmentation gene expression, melanin composition, and DNA repair responses, ultimately influencing the formation and persistence of dark cyclobutane pyrimidine dimers (CPDs) following ultraviolet (UV) exposure.

### Left panel

Ancestry-linked transcriptional variation (derived from GTEx skin transcriptomes) affects pigmentation and repair gene expression. Individuals with higher *SLC24A5* and reduced *TYR/TYRP1* expression (common in European ancestry) exhibit lighter pigmentation and reduced eumelanin synthesis. *XPC, DDB2*, and *POLH* (orange) represent the DNA repair system; *XPC* expression scales positively with *SLC24A5*, reflecting compensatory repair activation in lighter skin types.

### Middle panel

The melanin–chemiexcitation interface depicts epidermal architecture, where melanocytes transfer melanosomes containing eumelanin or pheomelanin to keratinocytes. Ultraviolet photons induce melanin oxidation and reactive oxygen species, producing delayed “dark” CPDs via chemiexcitation of DNA bases, hours after UV exposure. Eumelanin acts as an antioxidant and photoprotective filter, while pheomelanin generates ROS and triplet-state intermediates that propagate DNA damage (5, 6, 46).

### Right panel

Comparative outcomes highlight that lighter skin (European ancestry) experiences higher UV penetration, greater chemiexcitation, and elevated repair gene activation that only partially compensates for the increased damage load, resulting in higher dark CPD persistence and mutation risk. Darker skin (African ancestry) maintains stronger eumelanin protection, reduced oxidative stress, and lower demand on DNA repair pathways, yielding minimal dark CPD formation (1, 2, 55–57).

Together, the framework integrates evolutionary genomics, transcriptomic variability, and biophysical UV responses to explain ancestry-dependent differences in UV-induced DNA damage and repair dynamics.

### Data and Code Availability

All data derive from publicly available GTEx v9 expression and genotype releases (dbGaP phs000424.v9) (11, 21, 48).

## ACKNOWLEDGEMENTS

We thank the GTEx Consortium for open data sharing and acknowledge the foundational discoveries of melanin chemiexcitation by Premi and Brash (5, 6, 31, 32). Computational resources were provided by the institutional high-performance cluster.

## AUTHOR CONTRIBUTIONS

**Conceptualization** – Kingsley Essel Arthur and Christabel Martey;

**Data curation** – Kingsley Essel Arthur;

**Formal analysis** – Kingsley Essel Arthur;

**Methodology** – Kingsley Essel Arthur and Christabel Martey;

**Visualization** – Kingsley Essel Arthur and Christabel Martey;

**Writing – original draft** – Kingsley Essel Arthur

**Writing – review & editing** – Christabel Martey

## COMPETING INTERESTS

The authors declare no competing interests.

## ABBREVIATIONS

CI: Confidence Interval
CPD: Cyclobutane Pyrimidine Dimer
DCI: Dark-CPD Index
FDR: False Discovery Rate
GEO: Gene Expression Omnibus
GTEx: Genotype-Tissue Expression
GWAS: Genome-Wide Association Study
log2FC: Log2 Fold Change
NER: Nucleotide Excision Repair
PC: Principal Component
R^2: Coefficient of Determination
RMSE: Root-Mean-Square Error
RNA-seq: RNA Sequencing
ROS: Reactive Oxygen Species
RNS: Reactive Nitrogen Species
TPM: Transcripts Per Million
UV: Ultraviolet

## Notes

### Competing Interest Statement

The authors have declared no competing interest.

